# Evaluation of computational genotyping of Structural Variations for clinical diagnoses

**DOI:** 10.1101/558247

**Authors:** Varuna Chander, Richard A. Gibbs, Fritz J. Sedlazeck

## Abstract

**Background:** In recent years, Structural Variation (SV) has been identified as having a pivotal role in causing genetic disease. Nevertheless, the discovery of SVs based on short DNA sequence reads from next-generation DNA sequence methods is still error-prone, suffering from low sensitivity and high false discovery. These shortcomings can be partially overcome with the use of long reads, but the current expense precludes their application for routine clinical diagnostics. Structural Variation genotyping, on the other hand, offers cost-effective application as diagnostic tool in the clinic, with potentially no false positives and low occurrence of false negatives.

**Results:** We assess five state- of-the- art SV genotyping software methods that test short read sequence data. These methods are characterized based on their ability to genotype certain SV types and size ranges. Furthermore, we analyze their applicability to parse different VCF file sub-formats, or to rely on specific meta information that is not always at hand. We compare the SV genotyping methods across a range of simulated and real data including SVs that were not found with Illumina data alone. We assess their sensitivity and ability to filter out initial false discovery calls to assess their reliability.

**Conclusion:** Our results indicate that, although SV genotypers have superior performance to SV callers, there are performance limitations that suggest the need for further innovation.

## Background

With the continuous advancement of sequencing technologies, our understanding of the importance of Structural Variation (SV) is increasing[1]. Structural Variation has a critical role in evolution[2], genetic diseases (e.g. mendelian or Cancer) [2, 3] and the regulation of genes in different cells and tissues[4]. Furthermore, SVs compromise a substantial proportion of genomic differences between cell types, individuals, populations and species [1, 4-8]. Structural Variation is generally identified as being 50bp or longer genomic variations and categorized into five types: Insertions, Deletions, Duplications, Inversions and Translocations [9]. They are most often identified by leveraging paired-end, split read signals and coverage information[8].

Methods for the detection of SVs are still in their infancy, with some procedures reporting high (up to 89%) levels of false discovery[7, 8, 10-12] (i.e. SVs that are inferred due to artifacts, but not truly present in the sample) and between 10% to 70% false negatives[5, 7] (i.e. missing present SVs in the samples). Although the performance of these methods can improve by the use of long DNA sequence reads, this is often not practical due to high sequencing costs [13-15]. Therefore, using short reads alone significantly hinders SV discovery for routine clinical diagnosis [16].

An additional challenge is the interpretation of the possible functional consequences of SVs. Despite the availability of existing methods to compare SVs (e.g. SURVIVOR [5]) and to study the potential impact of SVs on genes (VCFanno [17], SURVIVOR_ant [18]), there is still a paucity of methods to assess their allele frequency among the human populations. These issues lead to problems that hinder routine screening for SVs in patient data and limit their proper recognition and characterization for clinical diagnoses.

The identification of SVs that have been previously identified in different samples is, in principle, easier than *de novo* detection. For known SVs it is possible to computationally detect SVs directly from short read DNA sequence data in individual per patient samples, guided by the expected position of split reads and discordant paired reads that can confirm breakpoints. This is less demanding than calling *de novo* SVs, since we focus only on specific genomic regions. This approach reduces the false discovery rates and therefore, reduces the potential misdiagnosis of patients. In addition, the false negative rate can be reduced as it is easier to genotype a variant than to identify a new SV. Genotyping known SVs has further the advantage that previous studies are likely to have annotated the event, and its possible association with certain diseases. Such events are recorded in SV databases (e.g. dbVar [19]).

In this paper, we review the current state of SVs genotyping methods and investigate their potential applicability for clinical diagnosis. In particular, we address whether these SV genotypers can re-identify SVs that short read callers were initially blind to (over GiaB [20] call sets) and how they perform for initially falsely inferred SVs. We precisely map out which genotypers operate on which types of SVs and their ability to genotype SVs based on sizes.

## Analyses

### Existing methods

Here we assess SVs genotyping methods: DELLY [21], Genome STRiP [22], STIX[23], SV2 [24] and SVTyper [25]. They share a common feature in which they require a bam file of the mapped reads and a VCF file that will be genotyped for SVs as inputs. **Table 1** lists their dependencies and their ability to genotype certain types of SVs.

**Table 1:**
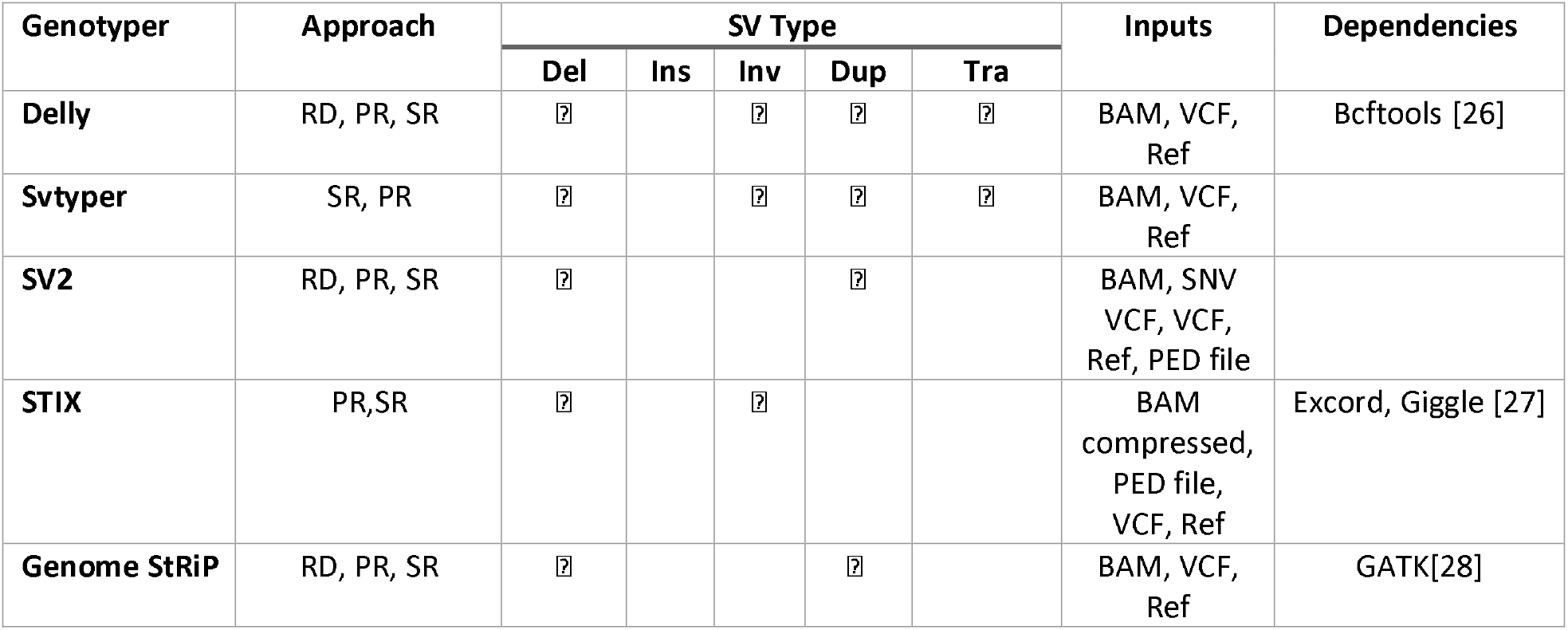
Overview of the SV genotypers assessed here and their ability to assess different SV types. RD: read depth, SR: split reads, PR: paired end reads

Overall, they can be divided into groups that support only two SV types (e.g. Genome STRiP) up to methods that support all SV types (SVTyper and DELLY), but require specific meta-information to do so. In the following, we give a brief description of each method that we assessed. Further insights can be obtained from their respective publications or manuals.

DELLY[21] is originally an SV caller that includes a genotype mode to redefine multi-sample VCFs. It operates on split and paired-end reads to genotype deletions, duplications, inversions and translations. However, for all types except the deletions, DELLY requires a sequence resolved call in its own format to be able to estimate the genotype.

Genome STRiP[22] genotypes only deletions and duplications. The unique aspect of Genome STRiP is that it was designed to genotype multiple samples simultaneously. It requires the GATK pipeline and prepackaged reference metadata bundles.

STIX[23], which is the most recently developed method included here, utilizes a reverse approach to the previous two examples. First, STIX extracts the discordant read pairs and split reads and generates a searchable index per sample. This index can then be queried if it supports a particular variant call. Noteworthy, STIX in the current form only provides information on how many reads support a variant rather than the genotype itself. This is done with a flag describing whether the reads are supported by a particular variant and the number of reads supporting it.

SVTyper[25] uses a Bayesian likelihood model that is based on discordant paired-end reads and split reads. It was designed to genotype deletions, duplications, inversions and translocations. For the latter, however, SVTyper requires specific ID tags provided by Lumpy[29] to complete genotyping.

SV2[24] uses a support vector machine learning to genotype deletions and duplications based on discordant paired-end, split read and coverage. Furthermore, it was the only SV genotyper assessed here that leverages SNP calls for its prediction.

### Evaluation of SVs computational genotypers based on simulated data

To first assess the performance of genotyping methods for SVs, we simulated data sets with 100bp Illumina like paired-end reads. Each data set includes 20 SVs simulated for a certain SVs type (duplications, indels, inversions and translocation) and a certain size range (100bp, 250bp, 500bp, 1kbp, 2kbp, 5kbp, 10kbp, 50kbp). For each of the data sets, we called SVs using SURVIVOR[5] based on a union set of DELLY, Manta[30], Lumpy[29] calls to include true positive as well as false positive SVs calls (see methods). Given the nature of the simulated data, we only observed 17 false discoveries while we were missing 17.25% of the simulated SVs. **Supplementary Table 1** shows the results for the discovery set over the 32 simulated data sets based on 640 simulated SVs on chr21 and chr22.

The generated VCF files were taken as input for five SV genotyper callers: DELLY, Genome STRiP, SV2, STIX and SVTyper. **Figure 1** provides an overview with respect to the ability to discover SVs in the first place (SURVIVOR). **Supplementary Table 1** shows the result for all genotypers applied to the 32 simulated data sets. Interestingly, we observed that not all methods accept a standardized VCF file and certain SV types require unique information. For example, while SVTyper is able to genotype deletions, inversion and duplications, it will just work on BND (translocation) events if the ID pairs provided by Lumpy are included in the VCF file. Also DELLY, which is also capable to infer deletions, inversion, duplications and translocations types of SVs is only able to genotype deletions given a standardized VCF without the extra information.

**Figure 1:**
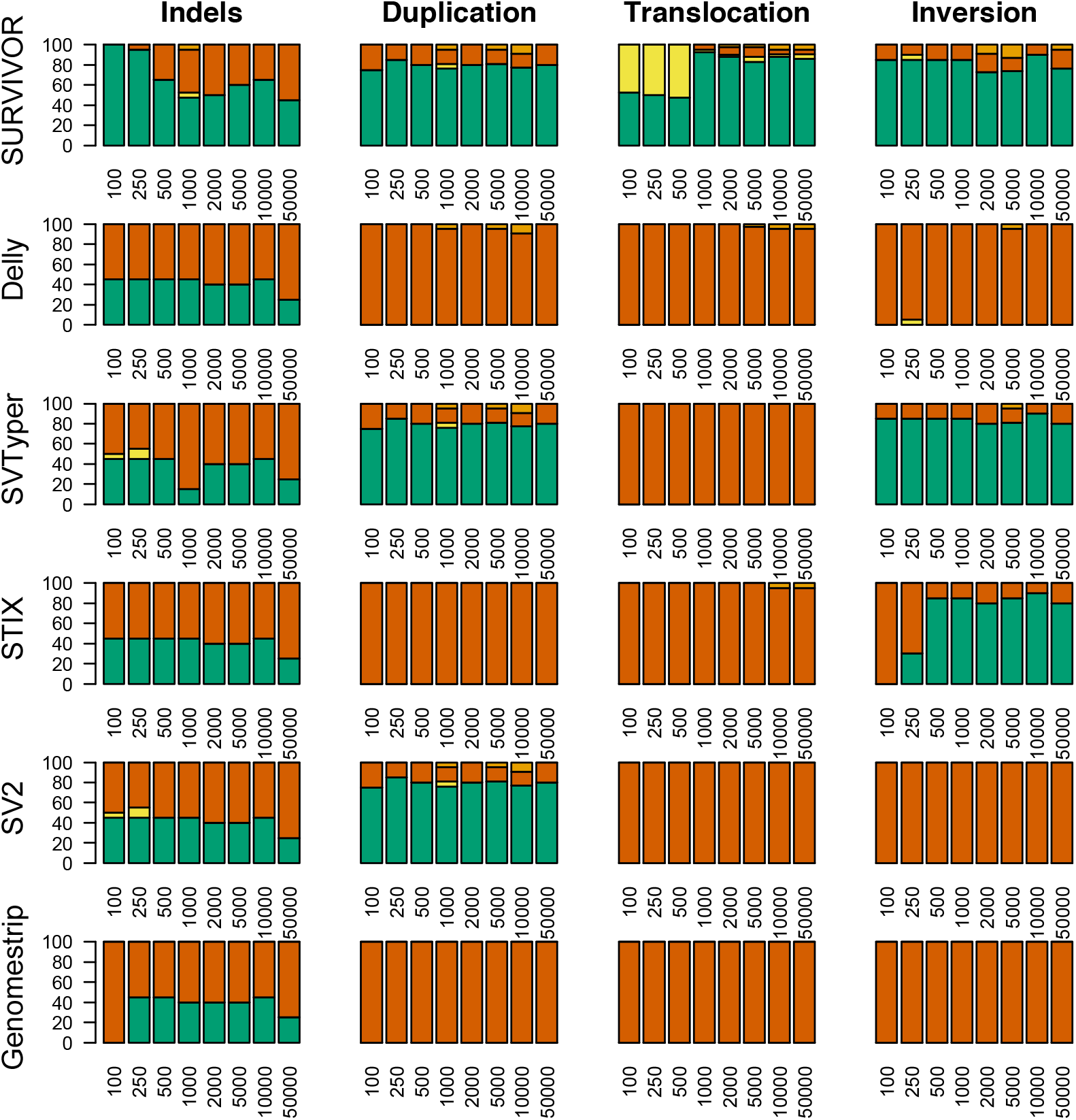
Evaluation of Illumina like reads to assess the SV genotyper ability to re-identify certain types and size ranges (x-axis). The colors indicate the ability of the SVs being detected/ genotyped either precisely (green), indicated (yellow), not detected (red) or falsely identified (brown) (see Methods). SURVIVOR is a union set of Delly, Lumpy and Manta to generate the VCF file as an input for the SV genotypers. The evaluation is based on the ability to discover SVs in the first place (SURVIVOR). Noteworthy, Delly and SVtyper can genotype more SVs given their costume information provided from their caller which are Delly and SVTyper, respectively.

The overall performance of each method was evaluated based on the input VCF generated by SURVIVOR. Thus, if all of the short read based SV callers were not able to resolve the insertions of 5kbp, then it would count as wrong/missed identified SV. In addition, we assessed the ability for the SV genotyper to filter out falsely called SVs. SVTyper (64.70%) had the highest rate of correctly genotyping SVs to be present, followed by SV2 (41.57%). Importantly, SV2 was able to genotype deletions and duplications, while SVTyper assessed deletions, duplications and inversions. Genome STRiP had the lowest (14.40%) success rate of all methods because it can only genotype deletions and duplications. This may reflect that Genome STRiP was designed primarily for population-based genotyping. SVTyper improved marginally (86.26%) when BND events, which represented translocations, were ignored, followed by the second best method SV2 (83.15%) when focused on deletions and duplications.

Next, we assessed the ability of the SV genotypers to reduce the false positives, i.e. initially wrongly inferred SVs. This will represent the scenario of accidentally genotyping a SV that is not represented in the sample due to sequencing or mapping biases. Over the 32 call sets, SURVIVOR had only 17 false positive calls for the simulated data. Genome STRiP performed best with filtering out all falsely detected SVs, but suffers from the lowest ability to genotype SV variations. STIX performed better as it can filter out 13 (81.82%) of the false positive SV calls. In contrast, STIX also achieved a higher (71.76%) performance for correctly identifying SVs. Although SVTyper had the highest accurately genotyped SVs it filtering out less of the false positives (69.70%) obtained during the discovery phase.

In summary, we observed that none of the methods is clearly superior for correctly genotyping and correctly filtering/non-reporting variations. Strikingly, none of the program were able to genotype insertions in the simulated data sets. Nevertheless, STIX and SV2 showed strong performance, with a good balance of sensitivity and being able to correctly discard false positives.

### Evaluation of SVs computational genotypers based on GiaB Ashkenazy Son

We further assessed genotyping of SVs calls based on the long read DNA sequence data from the Ashkenazy Son (HG002) reference sample. Specifically, we tested the currently released call set (v0.5.0) from GiaB, generated using sequence resolved calls from multiple technologies such as Illumina, PacBio, BioNano etc. and multiple SV callers and *de novo* assemblies based on these technologies, alone or in combination [20]. It is important to note that 8,195 of these SV calls could not be initially discovered with any Illumina assembly or caller but originated from PacBio based calls or BioNano based calling.

We are using this call set to genotype the SVs based on a 300x Illumina bam file for HG002 and compare the obtained SV genotype predictions to the genotypes reported by GiaB. The first observation was that most of the SV genotypers were unable to process the VCF file provided by GiaB. We used SURVIVOR to reduce the information included in the GiaB VCF file. Next, we filtered out the reported INS and complex events from this call set as most SV genotypers crashed while assessing these entries. Unfortunately, we were not able to run GenomeSTRiP successfully as it kept crashing even on a subset of these calls.

**Figure 2** displays the detectable deletions based on the GiaB call set (v0.5.0) per SV genotyper. STIX performed the best among all methods identifying 24,574 (78.74%) of the provided deletions. It is important to note that STIX does not currently report genotypes. Thus, we relied only on the information if STIX found reads that support this event rather than genotype information. DELLY performed as the second best identifying 18,528 (59.37%) deletions followed by SVTyper (34.24%) and SV2 (9.99%). Only 6.27% of the deletion calls from GiaB call set were genotyped by all SV genotype methods. Although this is a very low percent, it is positive that up to 78.74% of the deletions could be successfully identified out of 62,676 deletions (20bp+) in total. Noteworthy, 4921 deletions out of this set were never observed by any Illumina based caller or assembly. This highlights the potential benefit of using SV genotypers.

**Figure 2:**
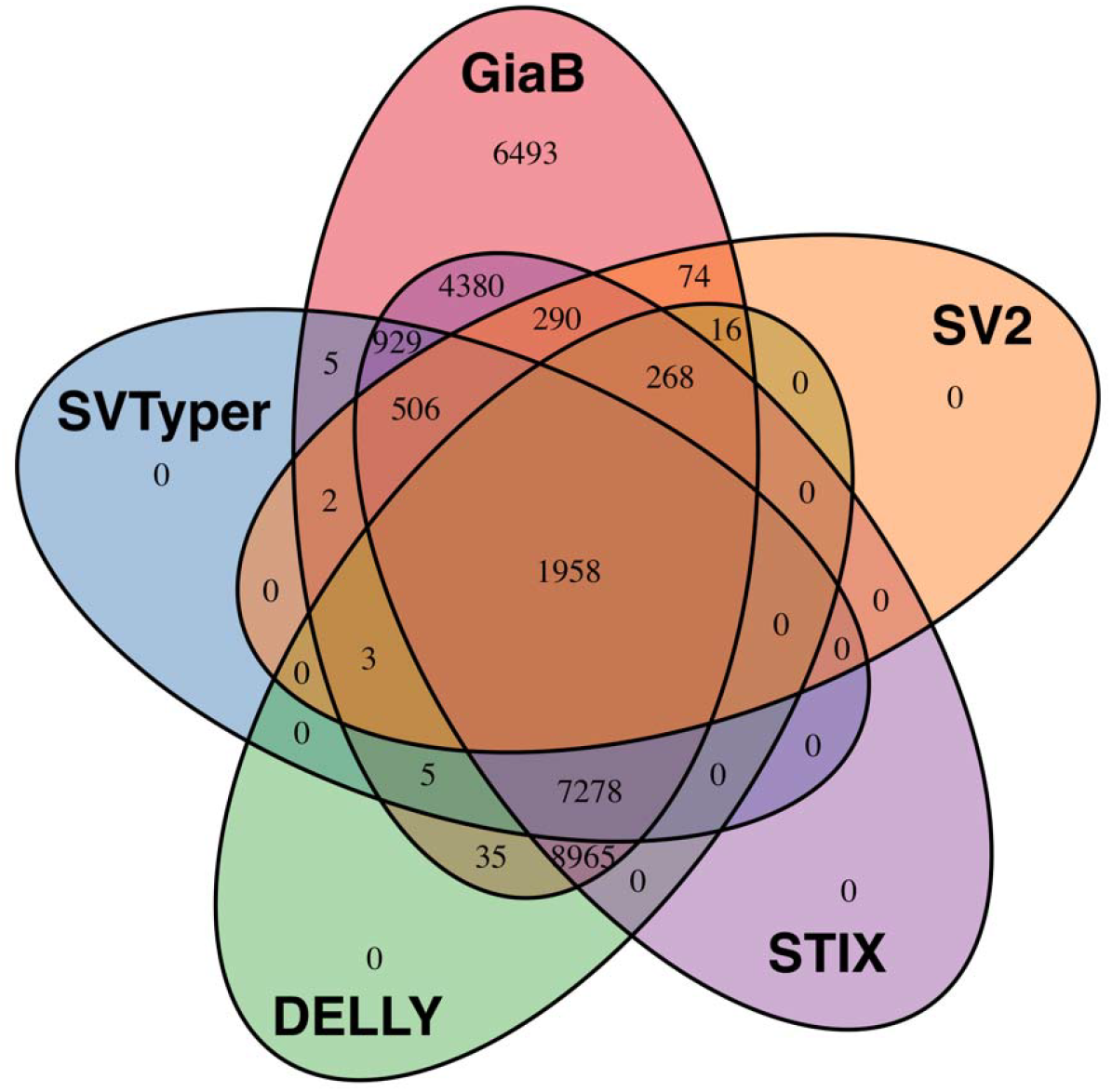
Evaluation based on GiaB call set v0.5.0 deletions only.

Next, we assessed the size regimes that SVs genotypers were able to recognize SVs. The deletions from GiaB call set 0.5.0 ranged from 20bp up to 997kbp with a median size of 36bp. All of the SV genotypers were able to identify deletions down to a size of 20bp. Interestingly we observed different median sizes of genotyped deletions, which represents the ability of specific methods to genotype small or large events. DELLY (31bp) had the lowest median SV size followed by SVTyper (32bp), STIX (35bp) and SV2 (116bp). Furthermore, DELLY (816kbp) genotyped also the longest SVs followed by STIX (694kbp), SV2 (656bkbp) and SVTyper (656bkbp).

When assessing the genotype concordance, DELLY performed the best with an agreement rate of 87.08% given that it identified the variant in the first place. SV2 achieved a 78.59% of genotype agreement, however it had one of the lowest recall rates (9.99%). SVTyper showed a 67.79% genotypes concordance. We did not evaluate STIX in this perspective since it does not report a genotype estimation in its current version.

In summary, STIX and DELLY performed the best in re-identifying the deletions reported by GiaB for HG0002. Furthermore, DELLY (87.08%) had also the highest agreement over the genotypes with the GiaB call set.

## Discussion

In this paper, we assessed the current state of SV genotyping methods. These methods are essential not just in assigning genotypes to discovery methods, but represents the ability to genotype already known, validated and functional annotated SVs. This latter aspect is important for clinical applications as it represents an early opportunity to reliably assess relevant SVs in the clinic.

Unfortunately, we discovered that many SV genotypers are only designed for their de novo SV caller counterpart in mind to report genotype estimations. This is especially obvious for DELLY, which can genotype all types subsequent to its discovery method, but only works on deletions based on a standardized VCF file. The same behavior, although reduced, can be observed by SVTyper that relies on specific IDs associated to translocations (in this case BND) events provided by Lumpy.

We were able to establish the first assessments of sensitivity and false discovery rate for SV genotypers not only focusing on Illumina detectable SVs, but further for SVs that were impossible to be discovered by Illumina alone. The latter becomes important as the field is continuously identifying SVs that based on long read technologies such as PacBio or Oxford Nanopore [13, 15]. These technologies often enable the detection of more complex SVs and also the detection of variations in regions that are hard to assess by Illumina alone. Thus, we rely on SV genotyper to assess if a particular SVs is present either in experiments with large sample sizes to evaluate the allele frequency or in clinical screenings to ease the diagnosis of patients [16].

Unfortunately, our study also highlights multiple general issues of SV genotyping methods. First, we observed that the methods tested here suffer from a limitation of SV types that they are able to assess. None of the methods were able to assess novel insertions that also represent repeat expansions, which is a subclass of SVs recognized to for their impact in cancer and other phenotypes. Second, most of the methods suffer from a very strict VCF formatting requirement ignoring the current standard and relied further on individual flags that are hard or impossible to regenerate.

Overall STIX performed well on simulated and GiaB based SVs calls. It showed a good balance of sensitivity vs false discovery and was able to be run on standard VCF files. Nevertheless, the lack of genotype estimations is a clear limitation since it is often relevant to know if a variation is heterozygous or homozygous. Overall, the current methods, although limited in performance, represents an advantage to diagnose patients with SVs compared to the discovery of SVs simply because of their much reduced false discovery rate.

## Potential implications

SVs genotyping represents a possibility to infer SVs in clinical diagnosis by solving the problems of false discovery and false negative called SVs, compared to the discovery SV methods. However, genotyping SVs methods seem to require additional development to improve their ability to operate on different size regimes and all types of SVs (including insertions). Here we presented an overview of the current state-of-the-art methods, which highlights the need to improve upon the current state of the art to enable SV diagnosis of patients in clinical setups.

## Methods

### Simulated datasets

We simulated 20 SVs per dataset each for a certain type (indel, inversions, duplication and translocation) and a certain size (100bp, 250bp, 500bp, 1kbp, 2kbp, 5kbp, 10kbp, 50kbp) for chr 21 and 22 using SURVIVOR simSV. These simulations included a 1% SNP rate. After the simulation of the sample genomes we simulated reads using Mason [31] with the following parameter “Illumina -II 500 -n 100 –N 39773784 -sq -mp -rn 2 “ to generate 100bp paired-end Illumina like reads. The reads were mapped with BWA MEM[32] using the –M option to mark duplicated reads to the entire genome (GRCh38-2.1.0). Subsequently, we ran Manta (v1.2.1), DELLY (v0.7.8) and Lumpy (v0.2.13) to call SVs over the simulated datasets. For each data set we generated a union call set based on all 3 callers using SURVIVOR merge (v1.0.3) allowing 1kbp distance and allowing only the same SV type to be merged. This union set, as well as the SV genotyper output, was evaluated with SURVIVOR eval for the following categories: Precise: calling an SVs within 10bp and inferring the correct type. Indicated: allowing a maximum of 1kbp between the simulated and the called breakpoints and ignoring the predicted type of SVs. Missing: a simulated SVs but not re identified. Additional: a SVs that was called, but not simulated. The results were summarized using a costume R script operating on the output of SURVIVOR available on request.

### SV genotyping: simulated data

For genotyping the simulated data set we used the union call VCF based on the SURVIVOR output as described above. We used DELLY (v0.7.8) specifying the output (-o), the vcf to be genotyped (-v) and the reference file (-g) as fasta and the bam file. We ran DELLY with the VCF file from SURVIVOR over the SV discovery caller. The obtained output from DELLY was converted using bcftools view (v1.7 (using htslib 1.7)) [26] to obtain a VCF file and was filtered to ignore genotyped calls with 0/0. SVTyper (v0.1.4) was used on the VCF generated from SURVIVOR based on the discovery phase. We filtered the obtained VCF for genotypes that could not have been accessed by SVTyper. SV2 (version 1.4.3) was run on the SURVIVOR generated VCF file for SVs genotyping but required also a SNV file. We generated this SNV file using Freebayes (v1.1.0-46-g8d2b3a0-dirty) [33] with the default parameters. The resulting SNV file from Freebayes was compressed and indexed by bgzip and tabix –p vcf [34], respectively. SV2 report their result in three folders (sv2_preprocessed, sv2_features and sv2_genotypes) from which we used the result reported in sv2_genotypes to benchmark the method. Genome STRIP(v2.00.1774) was used following the suggested parameters and the VCF file generated by SURVIVOR. STIX (early version available over GitHub on April 6^th^ 2018) was used to index the bam file using giggle (v0.6.3) [27], excord (v0.2.2) and samtools (v1.7) [26] following the suggested pipeline. Next, we run STIX with “-s 500” on the VCF files from SURVIVOR and ignoring output VCF entries with “STIX_ZERO=1”, which filters out entries where STIX does not find any evidence for the SV.

### SV genotyping: GiaB

We obtained the GiaB SV call set (v0.5.0) from ftp://ftp-trace.ncbi.nlm.nih.gov/giab/ftp/data/AshkenazimTrio/analysis/NIST_UnionSVs_12122017/. The SNV calls were taken from here ftp://ftp://ftp-trace.ncbi.nlm.nih.gov/giab/ftp/release/AshkenazimTrio/HG002_NA24385_son/latest/GRCh37/ and the corresponding bam file from ftp://ftp-trace.ncbi.nlm.nih.gov/giab/ftp/data/AshkenazimTrio/HG002_NA24385_son/NIST_HiSeq_HG002_Homogeneity10953946/NHGRI_Illumina300X_AJtrio_novoalign_bams/HG002.hs37d5.300x.b am. The SVs call set needed to be filtered and reduced for just one sample (HG002) using cat and SURVIVOR and was subsequently filtered for deletions only. We ran all SV genotyping methods like described above. Subsequently, we filtered the results for genotypes: 0/1 and 1/1 with the exception of STIX. STIX was filtered base on if it reports reads to support the SVs or not. This was necessary since STIX does currently not report genotypes. After filtering we merged all data sets together including the original VCF provided using SURVIVOR with a maximum distance of 10bp and requiring the same types. We analyzed these merged calls based on if the original call set reported a genotype to be heterozygous or homozygous alternative. The Venn diagram was generated based on the support vector reported by SURVIVOR and the R package Venn.diagram. The length of the SVs that were able to be genotyped were extracted using awk filtering for existing calls.

## Availability of data and materials

The data sets used in this study are available here ftp://ftp-trace.ncbi.nlm.nih.gov/giab/ftp/data/AshkenazimTrio/analysis/NIST_UnionSVs_12122017/ and from ftp://ftp-trace.ncbi.nlm.nih.gov/giab/ftp/data/AshkenazimTrio/HG002_NA24385_son/NIST_HiSeq_HG002_Homogeneity10953946/NHGRI_Illumina300X_AJtrio_novoalign_bams/HG002.hs37d5.300x.b am. The simulated data sets are available on request.

## Authors’ contributions

VC and FS performed the analysis. VC, FS and RG wrote the manuscript. FS and RG directed the project.

## Ethics approval and consent to participate

Not applicable

## Consent for publication

Not applicable

